# The quantitative contribution of different Photosystem II compartments to non-photochemical quenching in *Arabidopsis*

**DOI:** 10.1101/2021.10.17.463719

**Authors:** Lauren Nicol, Vincenzo Mascoli, Herbert van Amerongen, Roberta Croce

## Abstract

Excess excitation energy in the light-harvesting antenna of Photosystem II (PSII) can cause irreversible damage to the photosynthetic apparatus. In periods of high light intensity, a feedback mechanism known as non-photochemical quenching (NPQ), induces the formation of quenchers which can safely dissipate excess excitation energy as heat. Although quenchers have been identified in more than one compartment of the PSII supercomplex, there is currently no quantitative description of how much NPQ is occurring at each of these locations. Here, we perform time-resolved fluorescence measurements on WT and antenna mutants lacking LHCII (NoL) and all peripheral antenna (Ch1 and Ch1lhcb5). By combining the results with those of steady-state fluorescence experiments we are able to estimate the intrinsic rate of NPQ for each plant and each PSII compartment. It is concluded that 60-70% of quenching occurs in LHCII, 15-20% in the minor antenna and 15-20% in the PSII core.

## Introduction

Photosystem II (PSII) is the light-driven enzyme catalyzing the initial reactions of oxygenic photosynthesis - oxidation of water and reduction of plastoquinone (Q_A_). In the thylakoid membranes of plants, PSII is surrounded by an antenna of light-harvesting complexes (LHCs), which are densely packed with chlorophylls (Chls) and carotenoids. The pigment-protein matrix facilitates both efficient light absorption and excitation energy transfer to the reaction center (RC) where photochemistry takes place.

Upon absorption of a photon and promotion of a Chl to the excited state, photochemistry can occur within 100-300 ps if the PSII RC is open (i.e. Q_A_ is oxidized) (*1*). This is fast enough to largely outcompete excited-state decay pathways of fluorescence, internal conversion and intersystem crossing, resulting in a photochemical quantum yield of ∼0.8-0.9 (*2*). If the PSII RC is closed (i.e. Q_A_ is reduced), the Chl *a* excited-state lifetime increases to ∼2 ns (*3, 4*), increasing the probability of triplet formation via intersystem crossing. This situation can occur in high light intensities, when photon absorption exceeds the capacity of the electron transport chain. Chl triplets can then react with molecular oxygen to produce reactive oxygen species, which damage the photosynthetic apparatus and cause rapid photoinhibition (*5, 6*).

To prevent this outcome, a feedback mechanism induces the formation of quenchers - providing an alternative excited-state decay pathway which safely dissipates the excess excitation energy as heat. This process is known as non-photochemical quenching (NPQ) and in vascular plants, is triggered by an increase in lumen proton concentration and subsequent protonation of the integral membrane protein, PsbS (*7*–*9*). Low pH in the lumen also activates violaxanthin de-epoxidase, responsible for the conversion of violaxanthin to zeaxanthin (*10*). What happens next is largely unknown, but PsbS and zeaxanthin are thought to induce quenching sites in components of the PSII supercomplex either directly or indirectly (*11, 12*).

The dimeric PSII supercomplex can be broadly split into three compartments: PSII core, minor antenna and LHCII. The PSII core is a multi-subunit complex containing the RC and an inner antenna system coordinating 35 Chls *a* per monomer (*13*). The minor antenna consists of three monomeric LHCs, Lhcb4, Lhcb5 and Lhcb6 per PSII core monomer. Together they coordinate 38 Chls and have an important role in structurally and energetically connecting LHCII to the core (*1, 14, 15*). LHCII is a trimeric complex comprised of varying combinations of the three isoforms Lhcb1, Lhcb2, and Lhcb3 (*16*). Depending on light conditions, there can be 1-4 trimers per monomeric core, together coordinating up to 168 Chls (*2, 17*).

Quenching mechanisms have been proposed for each compartment of the PSII supercomplex. Quenching in LHCII is largely thought to occur via an aggregation-dependent mechanism, whereby PsbS and zeaxanthin induce the reorganization of PSII supercomplexes, and promote the formation of LHCII clusters (*18*–*20*). Both the interaction with PsbS and zeaxanthin, and the formation of clusters supposedly allow it to access the quenched state at pH values found in the thylakoid lumen (*11, 21, 22*). In the quenched state a large part of the excitation energy is transferred from chlorophylls to carotenoids, which quickly dissipate it (*23*–*26*). An independent quenching site has also been proposed to be present in the minor antenna (*27, 28*). The formation of a carotenoid cation radical observed in isolated minor antenna was suggested to be responsible for the quenching in these complexes (*29*–*32*). However, recent results have shown that chlorophyll to carotenoid excitation-energy transfer is also responsible for quenching in Lhcb4, suggesting that the quenching mechanism is the same in all LHCs (*25*). Quenching in the PSII core has also been proposed, although determination of the exact mechanism requires further investigation (*28, 33*–*35*).

For the most part, the various quenching mechanisms have been studied *in vitro*, and there are questions surrounding their physiological relevance, or to which extent they contribute to NPQ *in vivo*. Knockout mutants lacking entire PSII compartments therefore represent a valuable tool in answering these questions and the construction of a mutant lacking all minor antenna (NoM) was an important step in this regard (*14*). The mutant had WT levels of NPQ although altered induction kinetics, leading the authors to propose a multi-site model in which quenching in minor antenna and LHCII operate on different timescales (*28*). However, due to the poor energetic connection of LHCII to the core, it was difficult to identify if the changes in NPQ were due to a loss of quenching sites or other factors (*36*).

We have recently reported a mutant of *Arabidopsis thaliana* (*Arabidopsis*) lacking Lhcb1, Lhcb2, and therefore all LHCII trimers (NoL). While the mutant still contains Lhcb3, this isoform is unable to form homotrimers and accounts for a relatively small portion (∼10%) of LHCII under normal growth conditions (*37, 38*). These plants displayed a ∼60% reduction in NPQ indicating that LHCII trimers are the dominant site of quenching (*39*). The chlorina mutant (Ch1), displayed a similar reduction in NPQ (*39*). This mutant lacks chlorophyllide *a* oxygenase (CAO) required for the synthesis of chlorophyll *b* and therefore cannot stably accumulate functional Lhcb proteins. Lhcb5 is the only protein accumulated to a significant amount, however, it has been reported to be an apoprotein, not binding any pigments (*35*). It was therefore concluded that there was a significant secondary site of quenching in the PSII core and the minor antenna could only have a marginal contribution.

The study did not attempt to accurately quantify the contribution of each quenching site to NPQ because NPQ was measured via the quenching of steady-state fluorescence using pulse-amplitude-modulated (PAM) fluorometry. The value of NPQ in this instance is presumably the ratio of the rate of NPQ and the decay rate in the absence of NPQ and PSII photochemistry. However, this is only correct if there are no disconnected and unquenched antenna complexes. A direct comparison of the NPQ value between WT and antenna mutants is problematic for two additional reasons. First, the rate of NPQ is influenced by antenna size and second, it assumes the decay rate in the absence of NPQ and PSII photochemistry is the same in all plants. In order to make a quantitative comparison between plants it is necessary to directly compare the intrinsic rate of NPQ i.e. the rate at which excitation energy decays when it is located on the Chl that is being quenched (the quenching site). In the current study, we perform time-resolved fluorescence measurements on WT and the antenna mutants, NoL, and Ch1, as well as Ch1lhcb5, which additionally lacks the Lhcb5 apoprotein (*35*). We then use both steady-state and time-resolved data to access the intrinsic rates of NPQ for all plants, thus allowing a quantitative assignment of NPQ to each compartment of the PSII supercomplex.

## Results

### Steady-state fluorescence

The steady-state fluorescence intensity of a leaf is proportional to the fluorescence yield and strongly depends upon the yield of the other Chl excited state decay pathways (*40*). The minimal fluorescence of a dark-adapted leaf F_o_ corresponds to the ‘open state’ of PSII reaction centres, where Q_A_ is maximally oxidised. In this scenario, almost all of the absorbed energy is used in photochemistry, hence, the yield of fluorescence is small (Fig. 1a). The maximal fluorescence of a dark-adapted leaf F_m_, corresponds to the ‘closed state’ of PSII RC, where Q_A_ is maximally reduced. Here, photochemistry is blocked, therefore the yield of fluorescence increases (Fig. 1a). The maximal fluorescence of the light-adapted leaf F_m_’, also corresponds to the closed state of PSII RC, but here NPQ opens up an additional pathway for Chl excited state decay and the yield of fluorescence decreases (Fig. 1b).

**Fig. 1.**
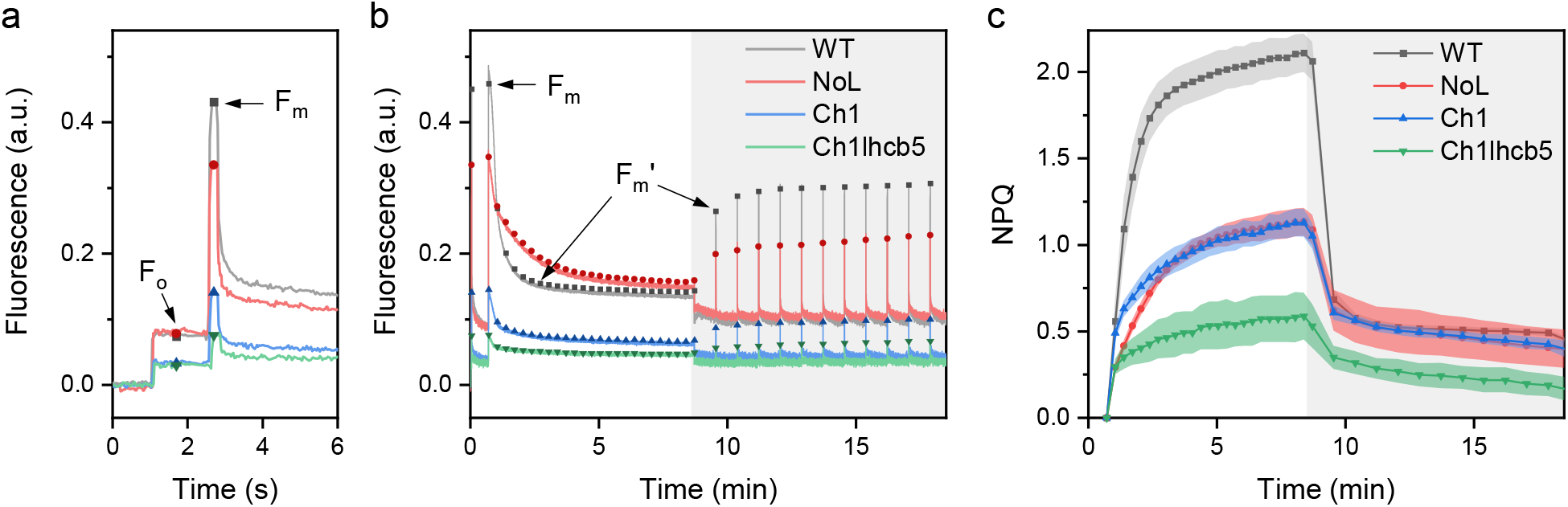
Steady-state fluorescence measurements of WT and antenna mutant leaves using PAM-fluorometry. **(A)** Raw fluorescence traces of dark-adapted leaves showing minimum chlorophyll fluorescence in the open state (F_o_) and maximum fluorescence in the closed state (F_m_), refer to 1b for the figure legend. **(B)** Raw fluorescence traces of chlorophyll fluorescence quenching upon illumination with 1,200 μmol photons m^-2^ s^-1^ for 8 minutes and chlorophyll fluorescence recovery in 10 minutes of darkness (grey background). F_m_ and examples of the maximum fluorescence in the light-adapted, closed state (F_m_’) are indicated. **(C)** Raw fluorescence converted to NPQ values. Data are mean ± s.d. (n = 3, 3, 6, 4 biological replicas for WT, NoL, Ch1 and Ch1lhcb5, respectively).

The NPQ value is calculated as the ratio between the difference of maximal fluorescence in the absence and presence of NPQ, and the maximal fluorescence in the presence of NPQ, i.e. (F_m_ − F_m_’)/F_m_’. Only the fast and reversible component of NPQ, called qE, is considered (see table 1). As shown in Fig. 1c and Table 1 all antenna mutants display a substantial reduction in NPQ compared to WT. A reduction in the NPQ value is often assumed to originate from a larger value of F_m_’, due to a smaller rate of quenching, however it could equally result from a reduction in F_m_, due to static quenching i.e. pre-existing quenching that does not require ΔpH for induction. The value of F_v_/F_m_, where F_v_ = F_m_ − F_o,_ is commonly used as an indicator of PSII quantum yield, and this value is also significantly reduced in all antenna mutants (Table 1). Likewise, this could indicate a decrease in F_m_ and/or an increase in F_o_, due to a disruption in the energetic connectivity of the PSII supercomplex.

**Table 1.** PAM parameters and pigment properties of WT and antenna mutants. Fv/Fm and NPQ data are mean ± s.d. (n = 3, 3, 6, 4 biological replicas for WT, NoL, Ch1 and Ch1lhcb5, respectively). NPQ values are reported as the difference in maximum level of NPQ in light and minimum level of NPQ following dark recovery, which corresponds to the fast and reversible NPQ component, also referred to as qE (*12*). Pigment data are are mean ± s.d. (n = 3 biological replicas).

| | $F_v/F_m$ | NPQ <sup>a,b</sup> | Chl $a/b$ <sup>c</sup> | Chl/car <sup>c</sup> |
| --- | --- | --- | --- | --- |
| WT | $0.83 \pm 0.00$ | $1.62 \pm 0.10$ | $3.31 \pm 0.02$ | $3.77 \pm 0.14$ |
| NoL | $0.77 \pm 0.01$ | $0.73 \pm 0.03$ | $4.55 \pm 0.15$ | $3.85 \pm 0.22$ |
| Ch1 | $0.76 \pm 0.02$ | $0.70 \pm 0.06$ | no Chl $b$ | $3.17 \pm 0.27$ |
| Ch1lhcb5 | $0.61 \pm 0.04$ | $0.40 \pm 0.09$ | no Chl $b$ | $3.10 \pm 0.32$ |

It is not possible to discriminate between these possibilities using steady-state fluorescence as the fluorescence intensity also depends on sample-specific parameters such as leaf area and Chl concentration. Instead, we perform time-resolved fluorescence measurements on samples in the various photosynthetic states. Here, average fluorescence lifetimes are directly proportional to fluorescence yield and independent of sample properties.

### Time-resolved fluorescence

Time-resolved fluorescence was used to determine both the excited-state lifetimes in the open and closed state (τ_o_ and τ_m,_ respectively) of WT and antenna mutants (Fig. 2). Average fluorescence lifetimes are directly proportional to fluorescence yields, and therefore provide “absolute” values of F_o_ and F_m_. An excitation wavelength of 468 nm was used to excite Chl *b* and carotenoids, both pigments quickly transferring excitation energy to Chl *a* (*1*). Fluorescence emission was detected at 680, 700 and 720 nm with the relative emission of PSI, which has a shorter average lifetime (<100 ps) (*41*) in comparison to PSII, increasing in the red. To obtain more quantitative information from the decay curves, they were globally analyzed and fitted to a sum of exponential decay components (Supplementary Table 1).

**Fig. 2.**
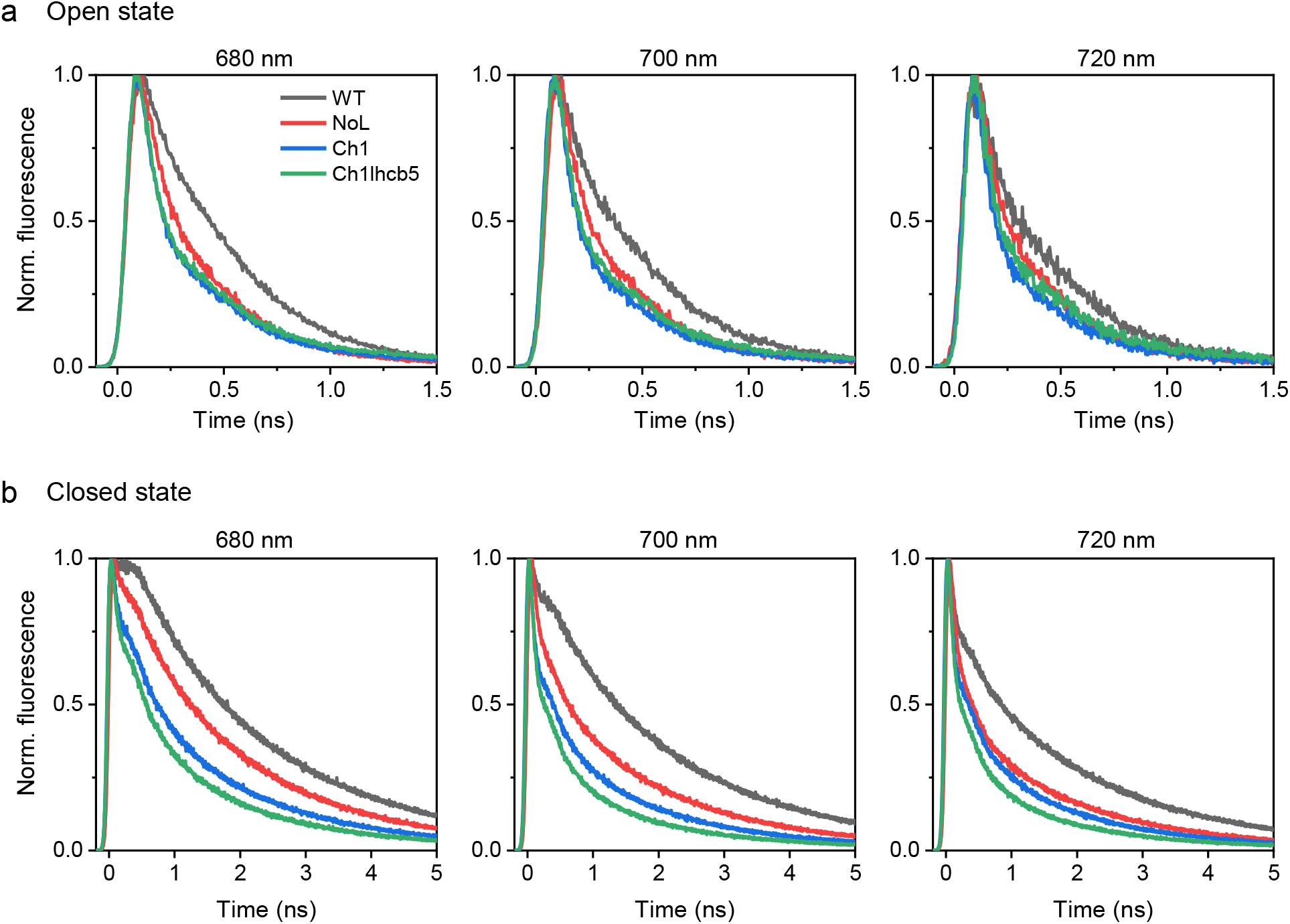
Normalized fluorescence decay curves of WT and antenna mutants. **(A)** Time-resolved fluorescence performed on thylakoid membranes in the open state. Traces are representative of 3 biologically independent replicas with similar results. **(B)** Time-resolved fluorescence performed on leaves in the closed state. The closed state is obtained via leaf infiltration with 200 μM DCMU. Traces are representative of 4, 7, 4, and 5 biologically independent replicas with similar results for WT, NoL, Ch1 and Ch1lhcb5, respectively. Samples were excited at 468 nm and fluorescence was detected at 680, 700 and 720 nm.

Open state measurements were performed on stacked thylakoid membranes. PSII was kept open throughout the measurement with low powered excitation and vigorous stirring of a large sample volume (Supplementary Fig. 1). The average fluorescence lifetimes of the mutants were shorter than that of the WT, in agreement with a smaller antenna size (Fig. 2a). A smaller antenna size reduces the distance an excitation has to travel to the RC and increases the probability of the excitation being located on the RC in an energetically equilibrated system. On the other hand, functionally disconnected antenna, closed PSII and free Chls all have long fluorescence lifetimes (> 1 ns), and the long-lived component present in all the mutants has a very small amplitude (≤ 4 %; Supplementary Table 1). Importantly, this indicates that the minor antenna complexes present in NoL are functionally connected to the PSII core.

Due to the difficulty in achieving a fully closed state in isolated membranes, these measurements were performed on leaves infiltrated with 3-(3,4-dichlorophenyl)-1,1-dimethylurea (DCMU), which blocks forward electron transport beyond Q_A_^-^. PSII was successfully closed with the excitation beam and measurements were performed under annihilation-free conditions (Supplementary Fig. 2). All antenna mutants display significantly shorter fluorescence lifetimes than WT (Fig. 2b). This static quenching lowers both steady-state F_v_/F_m_ and NPQ values in the antenna mutants, highlighting the need to calculate the rate of NPQ, which is independent of closed state lifetime.

### Intrinsic rate of NPQ

The overall quenching lifetime of NPQ *τ*_*NPQ*_ is the inverse of the rate of NPQ *k*_*NPQ*_ and can be determined using the acquired steady-state and time-resolved data with the equation

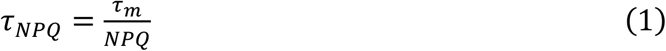

(see Supplementary Note 1 for details on how this equation is derived). The meaning of *τ*_*NPQ*_ can be easily understood by making a parallel with the trapping of excitations in the RC of PSII. It is well-known (see ref (*1*)) that the corresponding lifetime τ_*trap*_ of that process can be written as

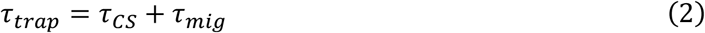

where the migration time τ_*mig*_ is the average time it takes for an excitation to reach the primary electron donor in the RC for the first time. It can also be considered as an equilibration time. The overall charge separation time τ_*CS*_ on the other hand denotes the charge separation after equilibration. This time is equal to the intrinsic charge separation time τ_*ics*_ (when the excitation is on the primary electron donor), divided by the probability *p*_*pd*_ that the primary donor *pd* is in the excited state according to the Boltzmann distribution

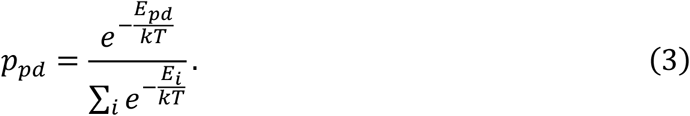

Here the summation runs over all pigments *i* with excited state *E*_*i*_ in the photosystem and includes the primary donor *pd* with excited-state energy *E*_*pd*_. The Boltzmann constant and absolute temperature are denoted with *k* and *T*, respectively. This then leads to

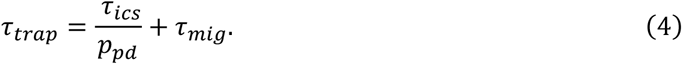

In case all *N* Chls *a* are isoenergetic

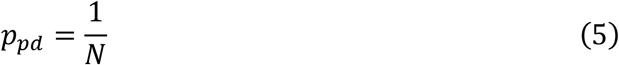

and

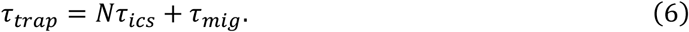

Exactly the same formalism can be used to describe quenching due to NPQ instead of quenching due to trapping in the RC:

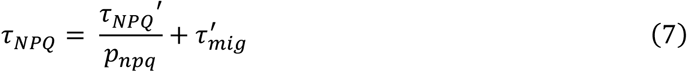

Here *τ*_*NPQ*_*’* is the inverse of the intrinsic rate of NPQ when the excitation is at the quenching site, *τ*_*mig*_*’* is the average time taken for an excitation to reach the quenching site and *p*_*npq*_ is the probability that the excitation is on the quenching site. In line with the Boltzmann distribution, *p*_*npq*_ can be approximated by 1/*N* where *N* is the number of isoenergetic Chl *a* molecules per supercomplex (Supplementary Table 2). If there is more than one quencher per supercomplex then the corresponding rates can simply be summed, leading to a combined overall rate of quenching. Given that both *τ*_*mig*_*’* and *p*_*npq*_ depend on antenna size, it is necessary to calculate *τ*_*NPQ*_*’* for a quantitative comparison of NPQ capacity between WT and the antenna mutants.

Like trapping in the RC, the nature of the quenching process lies somewhere in between a trap-limited case where *τ*_*mig*_*’* is infinitely short, and a migration-limited case where τ_*CS*_ is infinitely short. In the fully trap-limited case *τ*_*NPQ*_*’* can be calculated as:

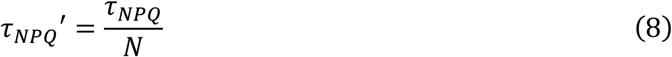

In the migration-limited scenario the exact value of *τ*_*NPQ*_ may depend on the location of the quenching site (see ref (*42*)) but here we estimate the upper limit for *τ*_*mig*_*’* to be equal to τ_*o*_. If trapping in the RC is fully migration-limited, τ_*o*_ is the average migration time from anywhere in the photosystem to the RC. This should be considered a safe estimate for *τ*_*mig*_*’* because energy transfer from the core antenna (CP43 and CP47) to the RC is relatively slow and the time to reach a quenching site is presumably somewhat shorter. Therefore, in the migration-limited scenario *τ*_*NPQ*_*’* can be calculated as:

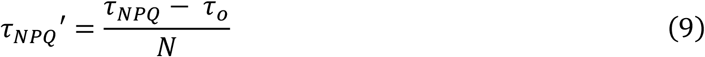

In the trap-limited scenario, WT, NoL, Ch1 and Ch1lhcb5 have intrinsic rates of quenching *k*_*NPQ*_*’* of (5 ps)^-1^, (13 ps) ^-1^, (20 ps) ^-1^ and (25 ps) ^-1^, while in the migration-limited scenario they have intrinsic rates of quenching *k*_*NPQ*_*’* of (3 ps)^-1^, (11 ps) ^-1^, (16 ps) ^-1^ and (21 ps) ^-1^. These rates are expressed relative to the rate in WT in Fig. 3a. Assuming that the quenching in the various PSII compartments occurs in a similar mechanistic way between WT and mutants, it is possible to calculate the *k*_*NPQ*_*’* for each PSII compartment according to:

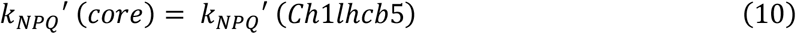

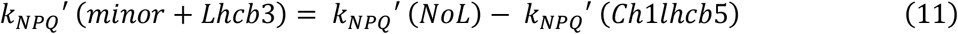

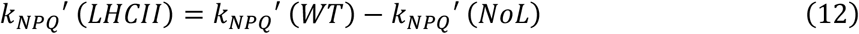

**Fig. 3.**
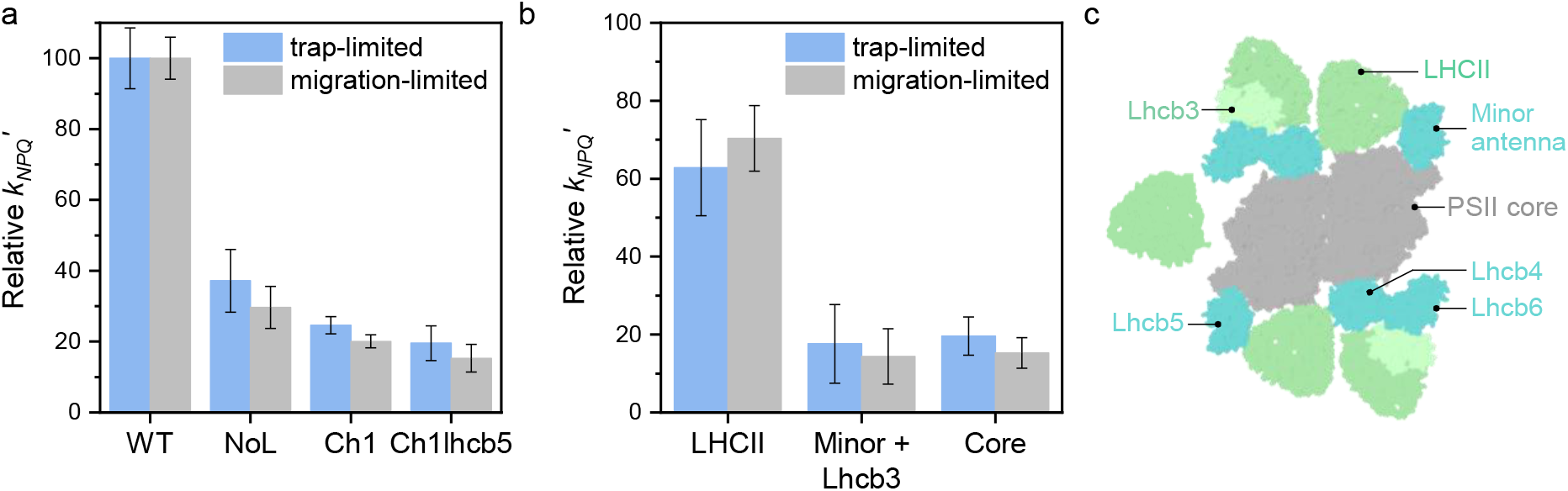
Intrinsic rates of NPQ (*k*_*NPQ*_*’*) in both trap- and migration-limited scenarios. **(A)** The *k*_*NPQ*_*’* of each antenna mutant relative to WT. Error bars represent standard deviation from at least three biological replicas. **(B)** The proportion of *k*_*NPQ*_*’* occurring in each PSII compartment in WT. Error bars represent standard deviation from at least three biological replicas. **(C)** Model of the dimeric PSII supercomplex, including an additional loosely bound trimer to represent the 2.5 LHCII trimers present per PSII core monomer in WT under normal growth conditions. PDB code: 5XNL (*15*).

The contribution of each compartment to NPQ can then be expressed as a proportion of the total *k*_*NPQ*_*’* occurring in WT. For example:

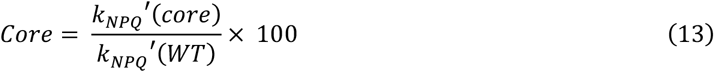

In both trap- and migration-limited scenarios this results in LHCII being responsible for 60-70% of quenching in WT, the minor antenna (together with Lhcb3) for 15-20% and the core for 15-20% (Fig. 3b).

Above we made the assumption that all Chl *a* molecules are isoenergetic, mainly for presenting a simplified calculation but this assumption influences the calculated quenching efficiency of the quencher by no more than 1% (see also SI).

At wavelengths above 700 nm PSI contributes substantially to fluorescence emission and so far, this contribution has been ignored. If the results are corrected for each plant’s individual PSI contribution, the absolute rates decrease, however the relative rates remain much the same (Supplementary Note 2; Supplementary Table 3-4; Supplementary Fig. 3), indicating that the conclusions are not affected by PSI.

## Discussion

Determining the location of NPQ in the PSII supercomplex is not trivial. Reverse genetic approaches, removing different combinations of pigment-protein complexes can often have unintended secondary consequences i.e. upregulation of other pigment-protein complexes (*43*) or poor connection of the remaining complexes to the PSII core (*14, 44*), making it difficult to accurately quantify the proportion of NPQ occurring in each compartment.

In our previous study, the level of NPQ was monitored in a mutant of *Arabidopsis* specifically deficient in LHCII (NoL), and the Chl *b*-deficient mutant (Ch1) in which almost all peripheral antenna complexes fail to accumulate (*39*). Lhcb5 is the only PSII antenna protein present in Ch1, however it is reported to be an apoprotein (*35*). Both mutants displayed a ∼60% reduction in NPQ, suggesting that the majority of quenching occurs in LHCII, a substantial amount of quenching occurs in the PSII core, and very little quenching occurs in the minor antenna. In the current study we have additionally measured the level of NPQ in the Ch1lhcb5 mutant, which also lacks Lhcb5 (*35*). Surprisingly, there is a large reduction in the level of NPQ in this mutant relative to Ch1, giving the impression that Lhcb5 is responsible for a significant amount of quenching. This is at odds with the single *lhcb5* knockout mutant, which is not affected in NPQ (*45*). The Chl1Lhcb5 mutant also displays a reduced value of F_v_/F_m_, as compared to the Ch1 mutant. This can be due to an increase in F_o_ or a reduction in F_m_; a reduction in the latter would cause an underestimation of NPQ and may explain the difference between Ch1 and Ch1lhcb5. To understand if this is the case and gain more insight into the steady-state fluorescence data we have performed time-resolved fluorescence on WT and the antenna mutants. In the open state, each sample displays a lifetime proportional to its antenna size, as has previously been observed in WT plants acclimated to different light conditions and for isolated PSII supercomplexes (*2, 46*). The near absence of long-lived fluorescence (>1 ns) in all antenna mutants suggests good functional connectivity of the remaining complexes. In NoL, this demonstrates that LHCII is not required for the functional connection of the minor antennae to the core, which is reasonable given that LHCII is the outermost antenna of the supercomplex.

Surprisingly, in the closed state, each plant displays a lifetime proportional to physical antenna size. The reduction of F_v_/F_m_ and, at least part of the reduction of NPQ in the antenna mutants can therefore be attributed to a reduction in F_m_ due to static quenching (i.e. quenching in the absence of ΔpH). The correlation of closed-state lifetime to physical antenna size suggests that LHCs stabilize the PSII core in a light-harvesting conformation, perhaps by controlling the macro-organization of complexes and preventing the clustering of PSII cores. It is also possible that the closed PSII reaction center is acting as a weak quencher. The quencher would become more populated in the absence of LHCs resulting in a shorter lifetime. It is worth noting that cyanobacterial PSII, which lacks integral membrane LHCs, also has a relatively short lifetime in the closed state *in vivo* (*47, 48*).

Using both steady-state and time-resolved fluorescence data, it was possible to calculate the *intrinsic* rate of NPQ (*k*_*NPQ*_*’*) for each plant. The small difference in *k*_*NPQ*_*’* between Ch1 and Ch1lhcb5 indicates that the stronger static quenching observed for Ch1lhcb5 largely accounts for the difference in steady-state NPQ values between Ch1 and Ch1lhcb5. The differences in *k*_*NPQ*_*’* between WT, NoL and Ch1lhcb5 provide us with the *k*_*NPQ*_*’* of each PSII compartment, and from this it is estimated that in WT plants ∼60-70% of quenching occurs in LHCII, ∼15-20% occurs in the minor antennae (and Lhcb3), and 15-20% occurs in the PSII core. If *k*_*NPQ*_*’* is normalized to the number of Chls *a* in each compartment to account for their differences in size, this results in LHCII being at least twice as efficient at quenching than either the minor antenna or the core. It has previously been reported that the rate of NPQ (or alternatively the number of quenchers) is higher in the presence of closed RCs than in the presence of open RCs (*49, 50*) which is in accordance with the presence of multiple quenching sites in the present study. The current NPQ experiments were performed in the presence of closed RCs and do not allow us to determine which quenching sites remain active for open RCs.

It should be emphasized that this model assumes that the loss of LHCII trimers in NoL or the loss of Lhcb complexes in Ch1lhcb5 does not affect the quenching properties of the remaining antenna or the core. This also implies that the quenching at all sites is independent from each other as has been previously suggested (*27, 28*).

In summary, the current study highlights the limitations of using steady-state fluorescence alone to assess the capacity of antenna mutants to perform NPQ. We show that relatively simple time-resolved measurements can be utilized to gain insight into the energetic connectivity of the PSII supercomplex, identify the presence of static quenching and estimate intrinsic rates of quenching. Our results demonstrate that NPQ does in fact occur in each compartment of the PSII supercomplex, providing support for previously proposed multi-site models of quenching.

## Methods

### *Arabidopsis* growth conditions and thylakoid isolation

*Arabidopsis thaliana* WT (Col-0), NoL, Ch1 and Ch1lhcb5 were grown under 120 μmol photons m^-2^s^-1^, 21 °C/18 °C, 16 h/8 h day/night cycle for 4–6 weeks. Thylakoid membranes were prepared as described previously (*51*) and could undergo one freeze-thaw cycle (flash frozen in liquid nitrogen, stored at -80 °C and thawed on ice) with no effect on results. The mutants used in this work have been extensively characterized previously (*35, 39, 52*). The pigment content, protein content, protein organization and spectra of the thylakoids of these plants are shown in Supplementary Fig. 4-6 and confirm the phenotypes.

### Pigment and protein analysis

Pigments were extracted from thylakoids membranes with 80% acetone. Room temperature absorption spectra of the pigment extracts were recorded on a Varian Cary 4000 UV–vis spectrophotometer. Chl *a*/*b* and Chl/Car ratios were calculated by fitting the absorption spectrum of the pigment extract with the spectra of the individual pigments (*53*). HPLC of the pigment extract was performed as described in ref (*53*) with modifications as reported in ref (*51*). Absorption spectra of thylakoids were measured on the Varian Cary 4000 UV–vis spectrophotometer equipped with an integrating sphere. SDS-PAGE electrophoresis was performed as in ref (*54*). Gels were loaded with 2.5 μg Chl and stained with Coomassie.

### PAM fluorometry

Using the modular Dual PAM-100 apparatus (Walz), whole leaves were excited with a measuring light (460 nm, 3 μmol photons m^−2^ s^−1^), and chlorophyll fluorescence emission was detected at wavelengths above 700 nm. The minimum level of fluorescence (F_o_) was detected following 1 hr of dark-adaptation. The maximum level of fluorescence (F_m_) was detected by applying a saturating pulse (635 nm, 180 ms, 12,000 μmol photons m^−2^ s^−1^) to dark-adapted leaves. Leaves were then light adapted for 8 min with an actinic light sufficient to saturate NPQ in all plants (635 nm, 1,200 μmol photons m^−2^ s^−1^) (*39*). This was followed by 10 min of recovery in darkness. The maximal level of fluorescence of the light adapted sample (F_m_’) was detected every 20 s with a saturating pulse. F_v_/F_m_ was calculated as (F_m_-F_o_) · F_m_^−1^. NPQ was calculated as (F_m_ − F_m_’) · F_m_’^−1^ and reported as the difference in maximum level of NPQ in light and minimum level of NPQ following dark recovery.

### Time correlated single photon counting (TCSPC)

Time-resolved fluorescence measurements were performed with a FluoTime 200 fluorometer (PicoQuant). Excitation was provided by a 468 nm laser diode and fluorescence emission was detected sequentially at 680, 700 and 720 nm. Thylakoid samples were diluted to an OD of 0.05 cm^-1^ at the Q_y_ maximum and a volume of 3 mL in 15 mM NaCl, 15 mM MgCl_2_ and 10 mM Hepes/KOH pH 7.5. The sample was kept at 20°C and stirred in a cuvette with a path length of 1 cm. The laser repetition rate was set to 10 MHz and the power reduced to 0.5-3 μW using neutral density filters (Supplementary Fig. 2). Each measurement had a maximum acquisition time of 15 min. A measurement with 680 nm detection was repeated at the end of each measuring sequence to ensure RCs were kept open throughout the measurement.

Whole leaves were taken from dark-adapted plants and incubated in a 0.3 M sorbitol, 10 mM Hepes pH 7.5, 200 μM DCMU solution for 4 hr. Successful DCMU infiltration was confirmed by ensuring that there was no difference in F_m_ under low actinic light illumination (i.e. 50 μmol photons m^−2^ s^−1^) and a saturating pulse (3,000 μmol photons m^−2^ s^−1^) in the Dual PAM. The leaf was inserted at an approximately 45-degree angle relative to both the excitation beam and the detector. Scattered light was blocked with an optical long-pass filter prior to detection. Laser repetition rate was set to 2.5 MHz and the power kept between 1-20 μW (Supplementary Fig. 3) to sufficiently close the PSII reaction centers while also avoiding singlet-singlet annihilation. A measurement with 680 nm detection was repeated at the end of each measuring sequence to ensure there was no photoinhibitory quenching occurring throughout the measurement.

Fluorescence decay curves were analyzed using FluoFit software (PicoQuant), following deconvolution of the instrument response function (IRF; 88 ps full width half maximum) measured from the ∼6 ps decay of pinacyanol iodide-dye dissolved in methanol (*55*). The data were globally fitted to multi-exponential decay functions with amplitudes A_i_ and associated fluorescence decay times *τ*_*i*_. The average fluorescence lifetimes were calculated according to *τ*_*avg*_ *= ΣA*_*i*_*\*τ*_*i*_*/ ΣA*_*i*._

## Author Contributions

Conceptualization: RC, HvA Investigation: LN, VM Supervision: RC

Writing—original draft: LN, HvA, RC Writing—review & editing: LN, VM, HvA, RC

## Data availability

All data needed to evaluate the conclusions in the paper are present in the paper and/or the Supplementary Materials.

