## Supplementary material for "The quantitative contribution of different Photosystem II compartments to non-photochemical quenching in *Arabidopsis*": SI

**This PDF file includes:**

Supplementary Text

Figs. S1 to S6

Tables S1 to S4

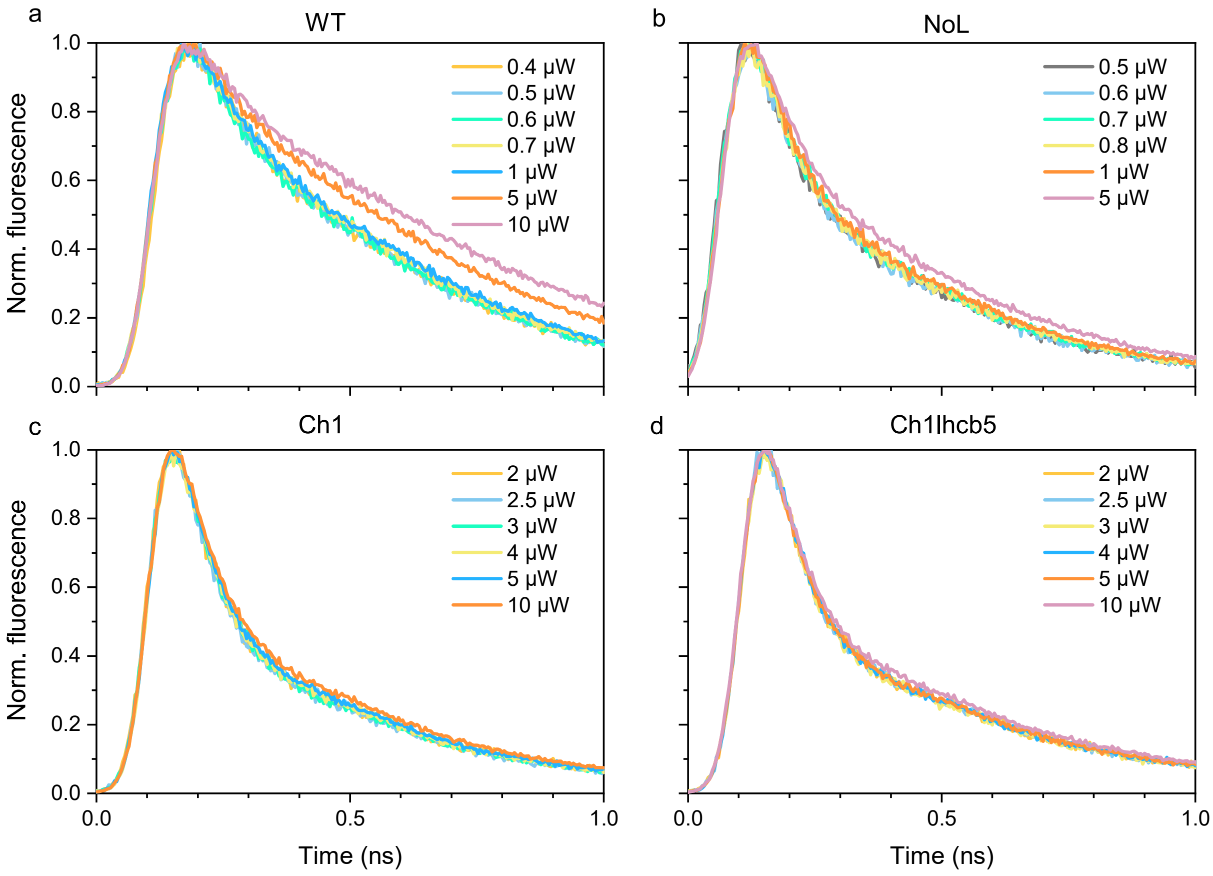


**Fig. S1. Power study to determine which excitation powers keep thylakoid membranes in the open state.** Samples were excited at 468 nm using different excitation powers and fluorescence was detected at 680 nm. Traces are representative of three biologically independent replicas with similar results. Powers of 0.5, 0.7, 3.0 and 3.0 μW were selected for further experiments on WT, NoL, Ch1, and Ch1lhcb5, respectively.

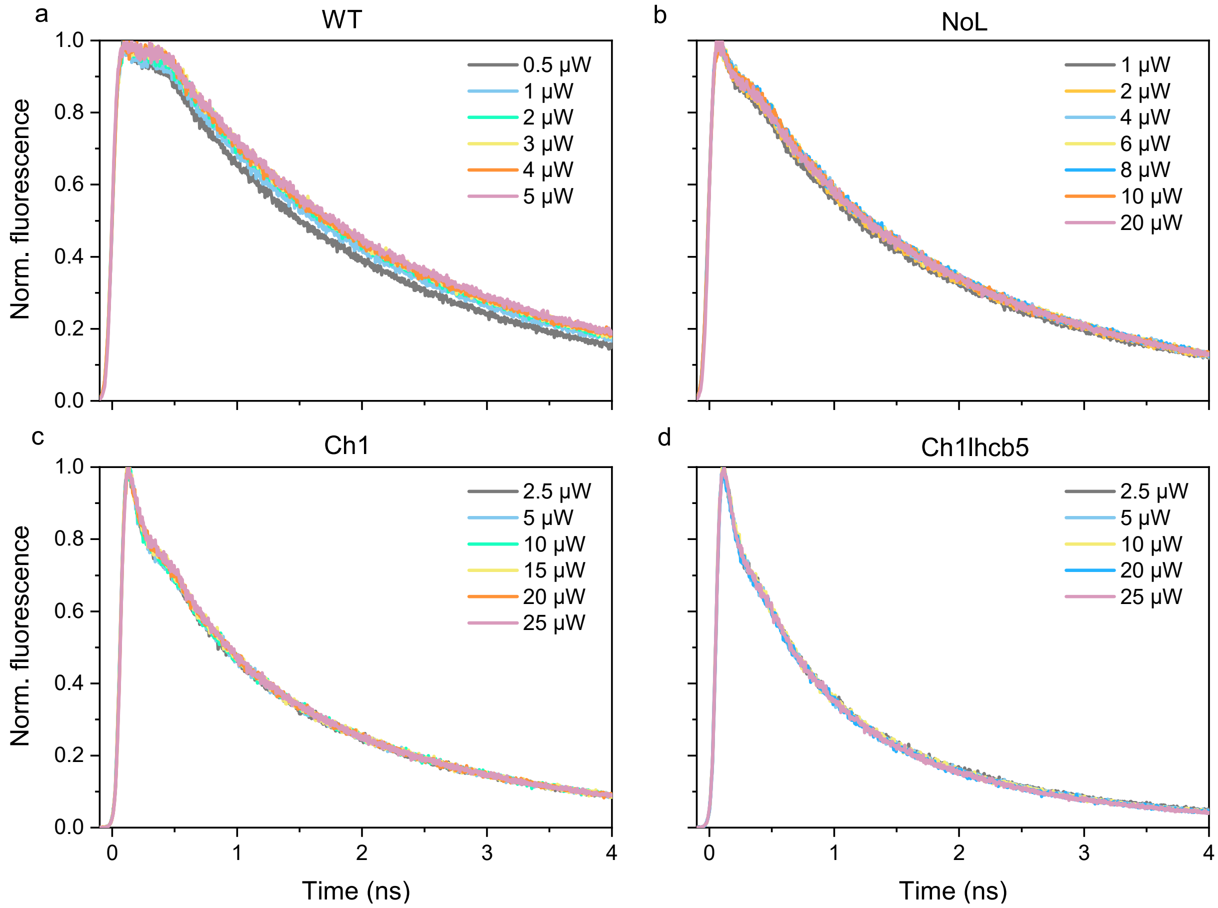


**Fig. S2. Power study to determine which excitation powers were sufficient to measure on DCMU-infiltrated leaves in fully closed state, while also avoiding singlet-singlet annihilation**. Samples were excited at 468 nm using different excitation powers and fluorescence was detected at 680 nm. Traces are representative of three biologically independent replicas with similar results. Powers of 2, 10, 20 and 20 µW were selected for further experiments on WT, NoL, Ch1, and Ch1lhcb5, respectively.

**Table S1. Global fitting of the fluorescence decay curves**[a]

**WT open state**

| | relative amplitudes (%) | | |
|---|---|---|---|
| **τ (ps)** | 680 nm | 700 nm | 720 nm |
| **107 ± 6** | 47 ± 6 | 59 ± 5 | 67 ± 5 |
| **352 ± 17** | 53 ± 6 | 41 ± 5 | 33 ± 5 |
| **$\tau_{avg}$ (ps)** | 236 ± 5 | 207 ± 7 | 188 ± 7 |

**WT closed state**

| | relative amplitudes (%) | | |
|---|---|---|---|
| **τ (ns)** | 680 nm | 700 nm | 720 nm |
| **0.09 ± 0.00** | 31 ± 5 | 49 ± 4 | 62 ± 2 |
| **1.05 ± 0.07** | 20 ± 4 | 17 ± 4 | 13 ± 3 |
| **2.39 ± 0.09** | 45 ± 4 | 34 ± 5 | 26 ± 3 |
| **$\tau_{avg}$ (ns)** | 1.4 ± 0.07 | 1.07 ± 0.07 | 0.82 ± 0.04 |

**NoL open state**

| | relative amplitudes (%) | | |
|---|---|---|---|
| **τ (ps)** | 680 nm | 700 nm | 720 nm |
| **102 ± 4** | 78 ± 2 | 79 ± 2 | 80 ± 2 |
| **341 ± 41** | 22 ± 2 | 21 ± 2 | 20 ± 2 |
| **$\tau_{avg}$ (ps)** | 155 ± 8 | 153 ± 9 | 150 ± 11 |

**NoL closed state**

| | relative amplitudes (%) | | |
|---|---|---|---|
| **τ (ns)** | 680 nm | 700 nm | 720 nm |
| **0.10 ± 0.00** | 36 ± 5 | 60 ± 7 | 62 ± 16 |
| **0.8 ± 0.04** | 18 ± 1 | 11 ± 2 | 10 ± 4 |
| **2.13 ± 0.10** | 40 ± 2 | 23 ± 2 | 19 ± 7 |
| **$\tau_{avg}$ (ns)** | 1.10 ± 0.06 | 0.68 ± 0.06 | 0.62 ± 0.24 |

**Ch1 open state**

| | relative amplitudes (%) | | |
|---|---|---|---|
| **τ (ps)** | 680 nm | 700 nm | 720 nm |
| **67 ± 3** | 75 ± 5 | 78 ± 4 | 77 ± 7 |
| **247 ± 10** | 22 ± 5 | 20 ± 4 | 20 ± 6 |
| **1101 ± 82** | 3 ± 1 | 3 ± 1 | 3 ± 1 |
| **$\tau_{avg}$ (ps)** | 133 ± 11 | 128 ± 11 | 131 ± 17 |

**Ch1 closed state**

| | relative amplitudes (%) | | |
|---|---|---|---|
| **τ (ns)** | 680 nm | 700 nm | 720 nm |
| **0.08 ± 0.00** | 51 ± 1 | 64 ± 1 | 70 ± 4 |
| **0.54 ± 0.02** | 23 ± 1 | 19 ± 1 | 17 ± 2 |
| **1.96 ± 0.09** | 27 ± 1 | 19 ± 1 | 16 ± 1 |
| **$\tau_{avg}$ (ns)** | 0.69 ± 0.02 | 0.52 ± 0.01 | 0.45 ± 0.03 |

**Ch1lhcb5 open state**

| | relative amplitudes (%) | | |
|---|---|---|---|
| **τ (ps)** | 680 nm | 700 nm | 720 nm |
| **64 ± 6** | 76 ± 4 | 76 ± 6 | 78 ± 6 |
| **231 ± 34** | 20 ± 2 | 20 ± 4 | 18 ± 4 |
| **1059 ± 108** | 4 ± 1 | 4 ± 2 | 3 ± 2 |
| **$\tau_{avg}$ (ps)** | 134 ± 26 | 136 ± 30 | 129 ± 30 |

**Ch1lhcb5 closed state**

| | relative amplitudes (%) | | |
|---|---|---|---|
| **τ (ns)** | 680 nm | 700 nm | 720 nm |
| **0.08 ± 0.00** | 55 ± 5 | 69 ± 9 | 73 ± 9 |
| **0.56 ± 0.04** | 25 ± 2 | 17 ± 1 | 14 ± 0 |
| **1.94 ± 0.12** | 19 ± 2 | 11 ± 1 | 9 ± 1 |
| **$\tau_{avg}$ (ns)** | 0.55 ± 0.06 | 0.37 ± 0.05 | 0.32 ± 0.04 |

[a] Samples were excited at 468 nm and fluorescence was detected at 680, 700 and 720 nm. The average fluorescence lifetime is calculated according to $\tau_{avg} = \Sigma A_i*\tau_i/\Sigma A_i$, where $A_i$ is the amplitude associated with $\tau_i$. Data in the open state are mean ± s.d (n = 3 biologically independent replicas). Data in the closed state are mean ± s.d (n = 4, 7, 4 and 5 biologically independent replicas for WT, NoL, Ch1 and Ch1lhcb5 respectively).

**Supplementary Text 1. Calculating the intrinsic rate of NPQ**

All Chl excited state decay pathways have an associated first-order rate constant (*k*) which is inversely proportional to the corresponding lifetime:

$k_f$ is the rate constant of fluorescence, also called the radiative decay rate.

$k_d$ is the rate constant of static quenching processes (internal conversion and intersystem crossing)

$k_{NPQ}$ is the rate constant of non-photochemical quenching

The yield of any excited state decay pathway is the ratio between its specific rate and the sum of the rate constants for all pathways of excitation decay. Therefore, the fluorescence yields of $F_m$ and $F_m'$ can be expressed in terms of kinetic parameters and substituted into the Stern-Volmer equation:

$F_m = k_f/(k_f + k_d)$ $F_{m'} = k_f/(k_f + k_d + k_{NPQ})$ $NPQ = (F_m - F_m')/F_m'$

$$NPQ = \frac{\frac{k_f}{k_f + k_d} - \frac{k_f}{k_f + k_d + k_{NPQ}}}{\frac{k_f}{k_f + k_d + k_{NPQ}}}$$

$$NPQ = \frac{k_{NPQ}}{(k_f + k_d)}$$

The outcome is that $k_{NPQ} = NPQ \cdot (k_f + k_d) = NPQ/\tau_m$ or, alternatively, $\tau_{NPQ} = \tau_m/NPQ$. In order to combine data from two different experimental set-ups (i.e. Dual Pam and TCSPC), they have to be obtained with similar excitation and detection wavelengths. Steady-state fluorescence comes from excitation at 460 nm and detection of wavelengths above 700 nm. For consistency, $\tau_m$ and $\tau_o$ are obtained with excitation at 468 nm and reflect the average of the fluorescence lifetimes detected at 700 and 720 nm.

In the main text we made the assumption that all Chl *a* molecules are isoenergetic, mainly for presenting a simplified calculation for the quenching efficiency of the quencher. Although the assumption is approximately correct (the Chl *a* absorption bandwidth is slightly larger than the thermal energy $k_BT$), it is not strictly required for obtaining the presented results. If the population probability of the quenched Chl *a* is for example twice as high as for the other pigments (because of a lower energy) then the calculated intrinsic rate of quenching would be half. However, the overall quenching efficiency by this pigment would essentially remain the same, because it is the product of the probability for the excitation to reside on the quencher (which is twice as high in this example) and the intrinsic quenching rate (which is approximately half as high). Although the product is not exactly identical, in all realistic cases it will not deviate by more than 1%.

**Table S2. Input values for the calculation of $k_{NPQ}$'.**

| | $\tau_o$ (ns)[a] | $\tau_m$ (ns)[a] | NPQ[a] | $k_{NPQ}$ ($ns^{-1}$)[b] | N Chl *a*[c] |
|---|---|---|---|---|---|
| WT | 0.20 ± 0.01 | 0.95 ± 0.05 | 1.62 ± 0.10 | 1.71 ± 0.15 | 120 |
| NoL | 0.15 ± 0.01 | 0.65 ± 0.15 | 0.73 ± 0.03 | 1.12 ± 0.27 | 68 |
| Ch1 | 0.13 ± 0.01 | 0.49 ± 0.02 | 0.70 ± 0.06 | 1.45 ± 0.14 | 35 |
| Ch1lhcb5 | 0.13 ± 0.03 | 0.35 ± 0.04 | 0.4 ± 0.09 | 1.15 ± 0.29 | 35 |

[a] Data are mean ± s.d (n ≥ 3 biologically independent replicas).

[b] Error represents standard deviation

[c] Based on known pigment and protein stoichiometries[2,13,15,35,39]

**Supplementary Text 2. Correcting for the contribution of PSI to fluorescence emission**

Fluorescence emission from leaves and thylakoid membranes originates from PSI and PSII. In the open state, both PSI and PSII are contributing to the fast component (~100 ps) of fluorescence decay. In the closed state, the lifetime of PSI remains approximately the same whereas the lifetime of PSII increases, meaning the fast component is primarily due to PSI. This contribution ($\tau_1$*$A_1$, averaged for detection at 700 and 720 nm) can be subtracted from $\tau_{o/m/m}$', allowing the extraction of PSII lifetimes (PSII $\tau_{o/m/m'}$), and the recalculation of NPQ based on quenching of PSII fluorescence only. For example:

$$\tau_m{}' = \frac{\tau_m}{NPQ + 1}$$

$$PSII\ \tau_{o/m/m}{}' = \frac{\tau_{o/m/m'} - (\tau_1 \times A_1)}{1 - A_1}$$

$$PSII\ NPQ = \frac{PSII\ \tau_m - PSII\ \tau_{m'}}{PSII\ \tau_{m'}}$$

Note that the contribution of PSI is similar in all plants (Supplementary Table 3) which correlates with the smaller PSI/PSII protein ratios observed in the antenna mutants (Supplementary Fig 3). Given that PSII core binds only 35 Chls and PSII-LHCII binds approximately 150 Chls per monomer, whereas the PSI core contains 100 Chls and PSI-LHCI contains 156 Chls – antenna mutants require a substantial (40-45%) reduction in the relative PSI protein content in order to maintain the excitation balance between photosystems.

**Table S3. Input values for the calculation of $k_{NPQ}$' corrected for PSI contribution.** Error represents standard deviation.

| | PSI ($\tau_1$*$A_1$) | PSII $\tau_o$ (ns) | PSII $\tau_m$ (ns) | PSII $\tau_m$' (ns) | PSII NPQ | PSII $k_{NPQ}$ ($ns^{-1}$) | N Chl *a* |
|---|---|---|---|---|---|---|---|
| WT | 0.050 | 0.33 ± 0.03 | 2.03 ± 0.12 | 0.71 ± 0.07 | 1.88 ± 0.19 | 0.93 ± 0.10 | 120 |
| NoL | 0.061 | 0.23 ± 0.03 | 1.50 ± 0.39 | 0.8 ± 0.23 | 0.87 ± 0.30 | 0.58 ± 0.24 | 68 |
| Ch1 | 0.054 | 0.23 ± 0.04 | 1.31 ± 0.07 | 0.7 ± 0.09 | 0.86 ± 0.11 | 0.66 ± 0.08 | 35 |
| Ch1lhcb5 | 0.057 | 0.26 ± 0.10 | 1.00 ± 0.15 | 0.66 ± 0.21 | 0.52 ± 0.18 | 0.52 ± 0.19 | 35 |

**Table S4. Comparison of the intrinsic rates of NPQ with and without PSI contribution.** Error represents standard deviation.

| | trap-limited $k_{NPQ}'$ ($ns^{-1}$) | | migration-limited $k_{NPQ}'$ ($ns^{-1}$) | |
|---|---|---|---|---|
| | **PSII** | **PSII + PSI** | **PSII** | **PSII + PSI** |
| WT | 111 ± 12 | 206 ± 18 | 161 ± 14 | 311 ± 18 |
| NoL | 39 ± 16 | 76 ± 18 | 45 ± 18 | 92 ± 18 |
| Ch1 | 23 ± 3 | 51 ± 5 | 27 ± 4 | 62 ± 6 |
| Ch1lhcb5 | 18 ± 7 | 40 ± 10 | 21 ± 10 | 47 ± 12 |

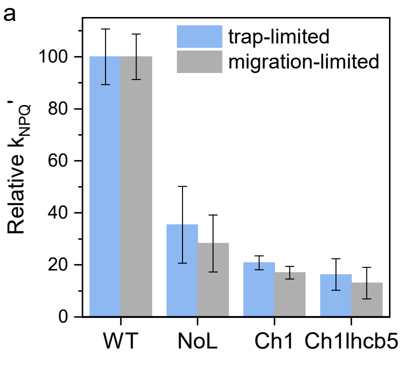


**Fig. S3. Intrinsic rates of NPQ corrected for PSI contribution.** (a) The $k_{NPQ}$′ of each antenna mutant relative to WT (b) the percentage of $k_{NPQ}$′ occurring in each compartment in WT. Error bars represent standard deviation.

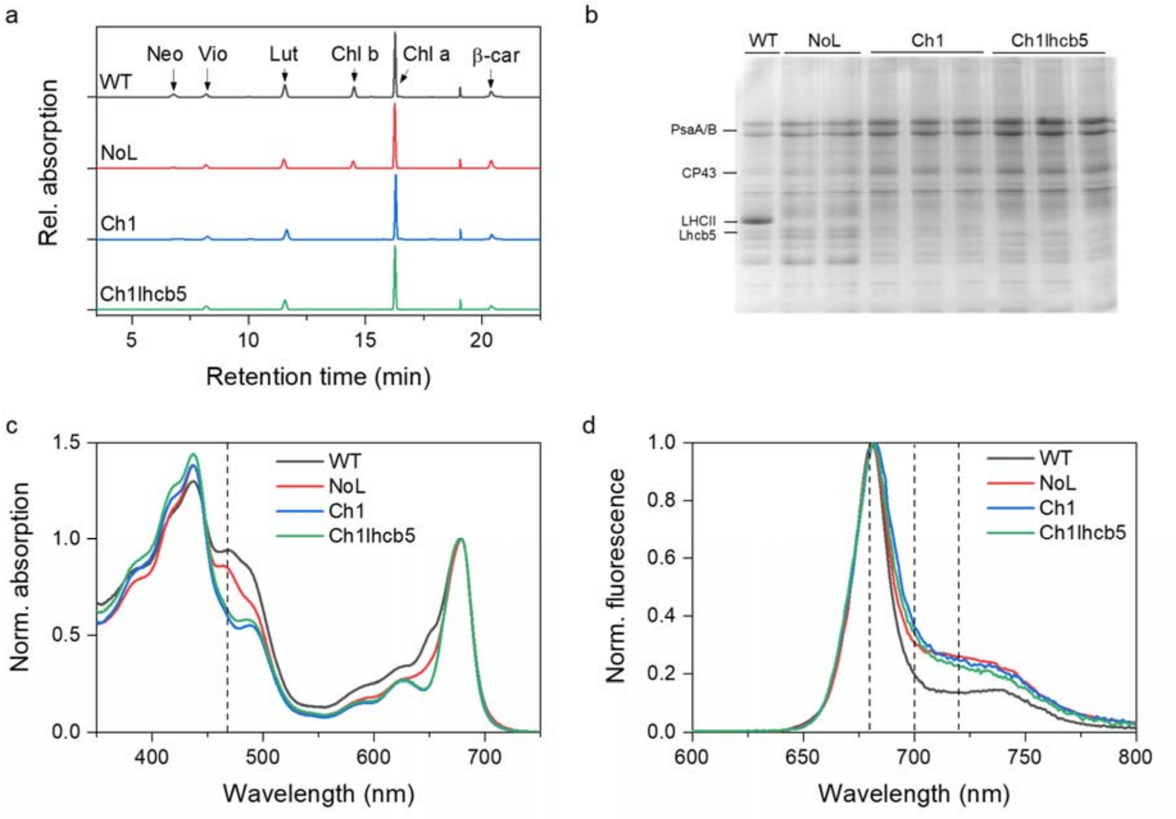


**Fig. S4. Pigment composition, protein composition, and spectral properties of WT and antenna mutant thylakoid membranes.** (a) Chromatographic profiles of the pigments extracted from thylakoid membranes normalized to the Chl a peak. Absorption is measured at 440 nm. Neo, neoxanthin; Vio, violaxanthin; Lut, lutein; β-car, β-carotene; Chl, chlorophyll. (b) Coomassie stained SDS-PAGE gel of thylakoids, with 2.5 µg Chl loaded per lane. (c) Absorption spectra of thylakoid membranes normalized to the $Q_y$ maximum. Dashed line indicates 468 nm. (d) Fluorescence spectra of thylakoid membranes normalized to the maximum value. Dashed lines indicate 680, 700 and 720 nm.

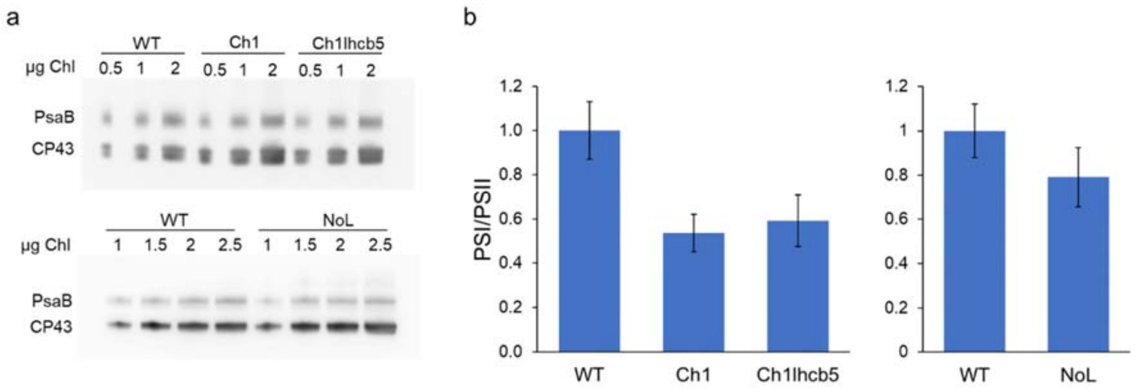


**Fig. S5. PSI/PSII protein ratios of WT and antenna mutants.** (a) Representative immunoblots of thylakoid membranes using antibodies against PsaB (PSI core subunit) and CP43 (PSII core subunit). Relative PSI/PSII protein ratios based on densitometric analysis of immunoblots. Data are mean ± s.d. (n = 4 biological replicas left panel, 2 biological replicas right panel).

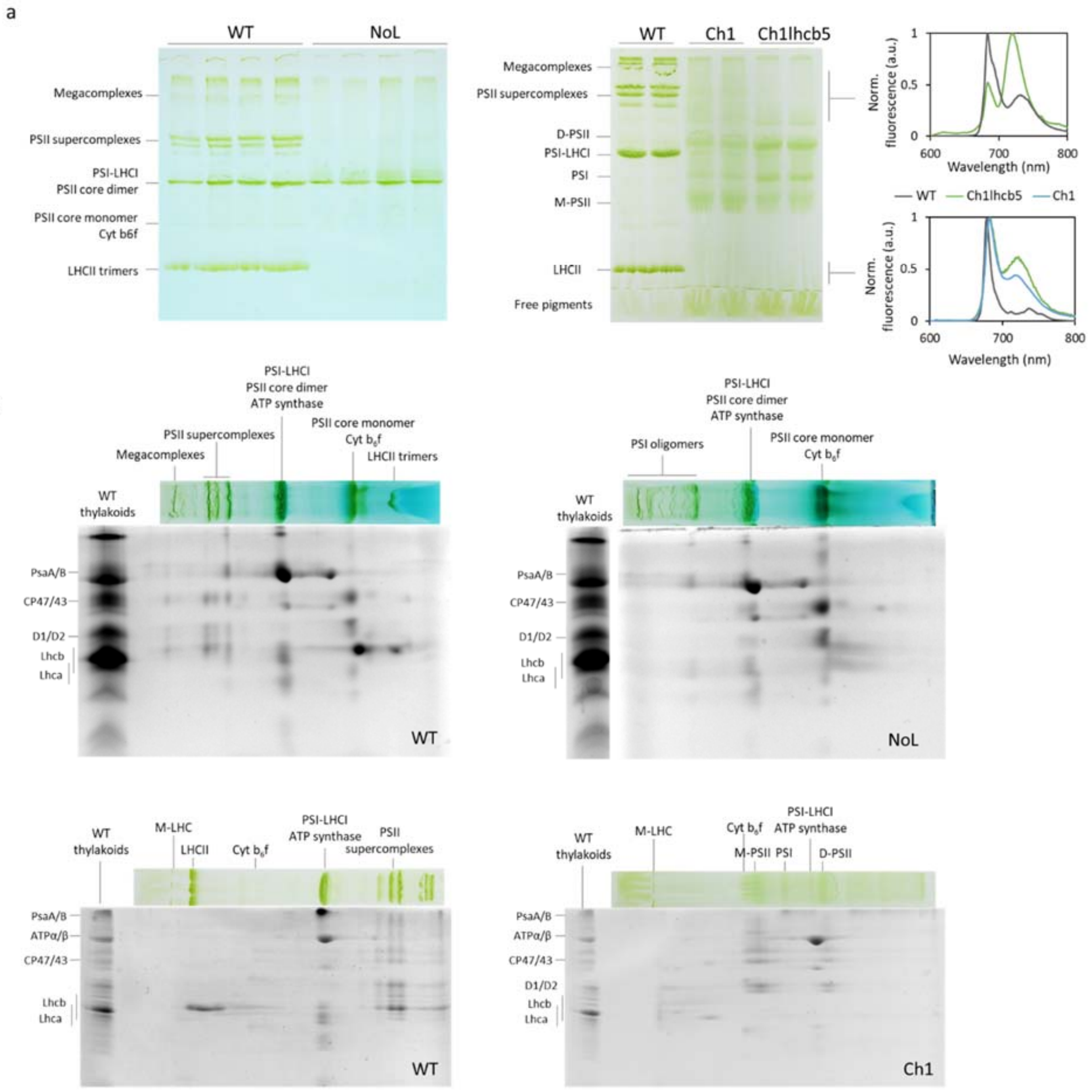


**Fig. S6 Protein organization in WT and antenna mutant thylakoid membranes. (a) Clear native-PAGE of WT, NoL,Ch1 and Ch1lhcb5 thylakoid membranes solubilized with 1% α-DM. Note the absence of PSII supercomplexes and LHCII in NoL.** In Ch1 and Ch1lhcb5, smeared bands present at the same height as PSII supercomplexes and LHCII give strong fluorescence emission at 720 nm. This indicates the presence of PSI oligomers and Lhcas, respectively. Two-dimensional gels of (c) NoL and (d) Ch1 further verifiy the absence of PSII supercomplexes and LHCII trimers in the antenna mutants.
